# Unlocking the biosynthetic potential of *Paenibacilli* through a genus-wide exploration of gene clusters for secondary metabolite production

**DOI:** 10.1101/2025.01.22.634348

**Authors:** Lijie Song, Matin Nuhamunada, Tilmann Weber, Ákos T. Kovács

## Abstract

The genus *Paenibacillus* is a prolific producer of secondary metabolites with diverse ecological and industrial applications. However, a comprehensive overview of the biosynthetic gene cluster (BGC) diversity and distribution throughout the genus has been limited. Here, we performed large-scale genome mining on 284 high-quality genomes and generated a non-redundant dataset of 126 representative genomes to explore the biosynthetic potential of this genus. A total of 3,273 BGCs were identified from the 284 genomes that clustered into 1,013 gene cluster families (GCFs), with 98.7% classified as unknown, indicating vast potential for novel secondary metabolite discovery in the *Paenibacillus* genus. Comparative analysis revealed significant phylogenetic and clade-specific distribution patterns of GCFs, with certain clades enriched in unique biosynthetic pathways while others exhibited low similarity to known BGCs, suggesting evolutionary adaptation to diverse ecological niches. This study uncovers the rich and largely untapped biosynthetic potential of the genus *Paenibacillus*, providing a foundation for future exploration of its natural products and their applications in biotechnology and medicine.

**IMPORTANCE:** Bacterial secondary metabolites have been instrumental in the development of antibiotics, antifungals, and other bioactive compounds. The genus *Paenibacillus* is an underexplored source of such metabolites, with significant potential for novel discoveries. By integrating genome mining and phylogenetic analysis, this study systematically characterizes the diversity, distribution, and novelty of biosynthetic gene cluster across the genus. The identification of clade-specific biosynthetic patterns and numerous unknown gene cluster families highlights *Paenibacillus* as a promising target for uncovering novel compounds with ecological and therapeutic relevance. These findings not only expand our understanding of bacterial secondary metabolite biosynthesis but also offer new opportunities for the development of sustainable biotechnological applications.

## INTRODUCTION

Secondary metabolites (SMs) are classically defined as groups of organic molecules that are not directly involved in the growth, development, or reproduction of producer organism, but that may confer ecological or physiological advantages unique to their producers (1). SMs have a wide range of applications in medicine, agriculture, and industry (2, 3). Bacteria are prominent SM producers offering a valuable source of bioactive compounds with potential therapeutic properties. These include antibiotics, antifungals, anticancer agents, and immunosuppressants (4). Since the discovery of streptomycin, chlortetracycline, nystatin, and numerous other antimicrobial compounds derived from *Streptomyces* in the 1940s to the present day, bacteria have contributed an enormous number of secondary metabolite compounds to the medical and pharmaceutical fields, as well as to scientific research (3).

The synthesis and production of secondary metabolites (SMs) in bacteria is frequently regulated by various genetic and environmental factors. In the case of microbes, the enzymes required for the formation of secondary metabolite compounds are typically encoded by a set of co-localized biosynthetic gene clusters (BGCs). These gene clusters contain all the genes necessary for the synthesis of specific secondary metabolites, including structural, regulatory, and transport-related genes (5). To better understand and utilize these secondary metabolites, scientists employ various bioinformatic tools and methods to identify and analysis these BGCs. One of the most widely employed bioinformatics tools in this field is antiSMASH, which has been specifically designed for the prediction and analysis of BGCs in bacterial and fungal genomes (6). BGCFlow is a powerful workflow that integrates antiSMASH with additional tools for data visualization, clustering, and functional annotation, rendering it particularly effective for large-scale comparative studies (7).

The utilization of these tools and connected databases (e.g. MIBiG, antiSMASH-DB, BIG-Fam) (8–10), particularly in conjunction with the advancement of genome sequencing studies and the availability of large-scale datasets, has enabled scientists to efficiently detect BGCs and infer SM products from a diverse range of bacterial species (11–14). A lot of bacteria have been identified that possess a high density of BGCs and secondary metabolites. Among these, the *Actinomycetes* and *Bacilli* have attracted particular attention due to the abundance of their secondary metabolites, especially of antibiotics and plant-promoting active compounds (15–21).

*Paenibacillus*, originally classified within the *Bacillus* genus but later reclassified as a separate genus in 1993 (22), is known to produce a wide range of biologically active secondary metabolites. According to the LPSN database (https://lpsn.dsmz.de), the total number of validly published names (including synonyms) for the *Paenibacillus* genus is 320. The majority of *Paenibacillus* are found in soil and frequently associated with plants, where these rhizosphere bacteria have been demonstrated to promote plant growth through a range of metabolic capabilities, including biological nitrogen fixation, phosphate dissolution, and the production of plant hormones (23–25). Another aspect of *Paenibacillus* that has become more notable in recent years is its ability to produce antimicrobial compounds. This has led to the genus becoming a focus of research connected to its potential as a biocontrol agent and natural antibiotic producer. For example, polymyxin, a known antibiotic produced by *Paenibacillus polymyxa*, is used to treat infections caused by Gram-negative bacteria. Other antimicrobial compounds produced by *Paenibacillus spp* include paenibacillin, polyxin, and lantibiotics, among others (26, 27). *P. polymyxa* E681, the representative and most studied strain of the species has been extensively analyzed for its BGCs, which are responsible for the production of at least six antibiotics, including polymyxin, fusaricidin, tridecaptin, paenilipoheptin, paenilan, and bacillaene-like antimicrobials (28), Pranav *et al.* identified 104 BGCs from the genome sequences of five *Paenibacillus alvei* strains, and BGCs encoding paenibactin, fusaricidin, and paenibacterin were found to be present in all five strains (29). Lebedeva *et al.* investigated the biosynthetic potential of two *Paenibacillus sp*. strains isolated from caves and revealed 21 and 19 BGCs in those strains (30). Kim *et al*. analyzed 89 genomes of *Paenibacillus* using antiSMASH, and the result revealed a total of 848 BGCs, with a significant majority (716 or 84.4%) classified as unknown, indicating a vast unexplored potential for new antibiotics within these species (31). Despite extensive research on the biosynthetic gene clusters (BGCs) and bioactive compounds of various *Paenibacillus* species, an up-to-date overview of the BGC composition and distribution across this genus remains elusive. Here, we report the analysis of 284 genomes of the *Paenibacillus* genus and demonstrate conserved and phylogenetically distributed BGCs that further support the potential of this genus for discovery of novel bioactive metabolites.

## RESULTS AND DISCUSSION

### Genome mining reveals the biosynthetic potential of the *Paenibacillus* genus

In this study, to establish a systematic insight on the BGC composition of the *Paenibacillus* genus, we obtained all available accession IDs for *Paenibacillus* genomes from the NCBI database as of 10 October 2024, and there were 1,969 available genome sequences in total. Since the accuracy and completeness of BGC identification are dependent on genome quality (12), the genomes were filtered according to their assembly level, completeness, and contamination, following the strategy described in the method section. This process yielded 284 genomes that were annotated by GenBank and deemed to be of high quality, and which were subsequently included in the complete dataset for this study (see Table S1). The genome size of the 284 genomes ranged from 3.97 M to 9.08 M, with 89.08% of them falling within the 5-8 M range. The GC % varied from 40% to 63.5%, with 96.48% of them falling within the 40-55% range (see Fig. S1 and Table S1). The GTDB-Tk (version 2.4.0) was employed for taxonomic assignment at the species level using GTDB release 09-RS220 (32, 33), resulting in a total of 266 genomes assigned to one of the 122 species identified. Of these, 89 were represented by a single genome, while some species have numerous genomes, including *P. polymyxa_B* (27 genomes), *P. odorifer* (22 genomes), and *P. polymyxa* (19 genomes). Additionally, 18 genomes lacked species assignments and were treated as unclassified species (see Table S1).

Utilizing antiSMASH v7.1.0 (6), a total of 3,273 BGCs were predicted from the 284 genomes, with an average of 11.25 BGCs per genome, ranged from 2 to 22 (see Fig. S2 and Table S2). The number of BGCs identified per genome differed significantly between species. For instance, the 9 *Paenibacillus_H larvae* genomes in this dataset had an average of 17.33 BGCs per genome, whereas the 22 *Paenibacillus odorifer* genomes had an average of only 5.73. In a previous study that was based on 89 genomes of *Paenibacillus* available in February 2020, Kim *et al*. obtained an average of 9.5 BGCs per strain (31). The higher number of BGCs observed in our dataset may be attributed to the higher percentage of *Paenibacillus polymyxa* and related species, which are BGC-rich species. No significant correlation was found between genome size and the number of BGCs per genome in this *Paenibacillus* dataset (see Fig. S3). This finding differs from those of other researchers in the field of actinomycetes, who observed a positive trend between the number of BGCs and genome size (34). It is noteworthy that only 16 of the 3,273 BGCs are situated at contig edges, indicating that 99% (3,257/3,273) of the BGCs are complete, and this observation supports the high quality of the dataset for this study (35).

### High diversity and novelty of BGCs of *Paenibacillus* species

The 3,273 BGCs identified in this dataset encompass the majority of all BGC types (Fig. 1a and Table S2). In summary, Non-ribosomal peptide synthetases (NRPS) and PKS-NRPS hybrids make up the majority of all BGCs with 435 and 413 respectively, representing 13.29% and 12.62% of the total BGCs. This finding is consistent with the results of previous studies on *Paenibacillus*. The third and fourth most abundant BGC types are cyclic lactone inducers and proteusins, which both belong to ribosomally synthesized and post-translationally modified peptides. There is a paucity of comprehensive data regarding the proportion of these BGC types in other published papers, likely due to the recent incorporation of annotations for these BGC types in the more recent version of antiSMASH (V.7.1.0). All genomes contain a variety of BGCs belonging to different types. The most widely distributed BGC is proteusins, which was identified in 234 out of the 284 genomes (82.39%). Other types of BGCs that are widely distributed across all 284 genomes (e.g. more than 50% of the genomes) include cyclic-lactone-autoinducer (190 genomes, 66.90%), lassopeptide (183 genomes, 64.44%), terpene (180 genomes, 63.38%), PKS-NRPS hybrids (175 genomes, 61.62%), and NRPSs (158 genomes, 55.63%) (Fig. 1b).

**Fig. 1.**
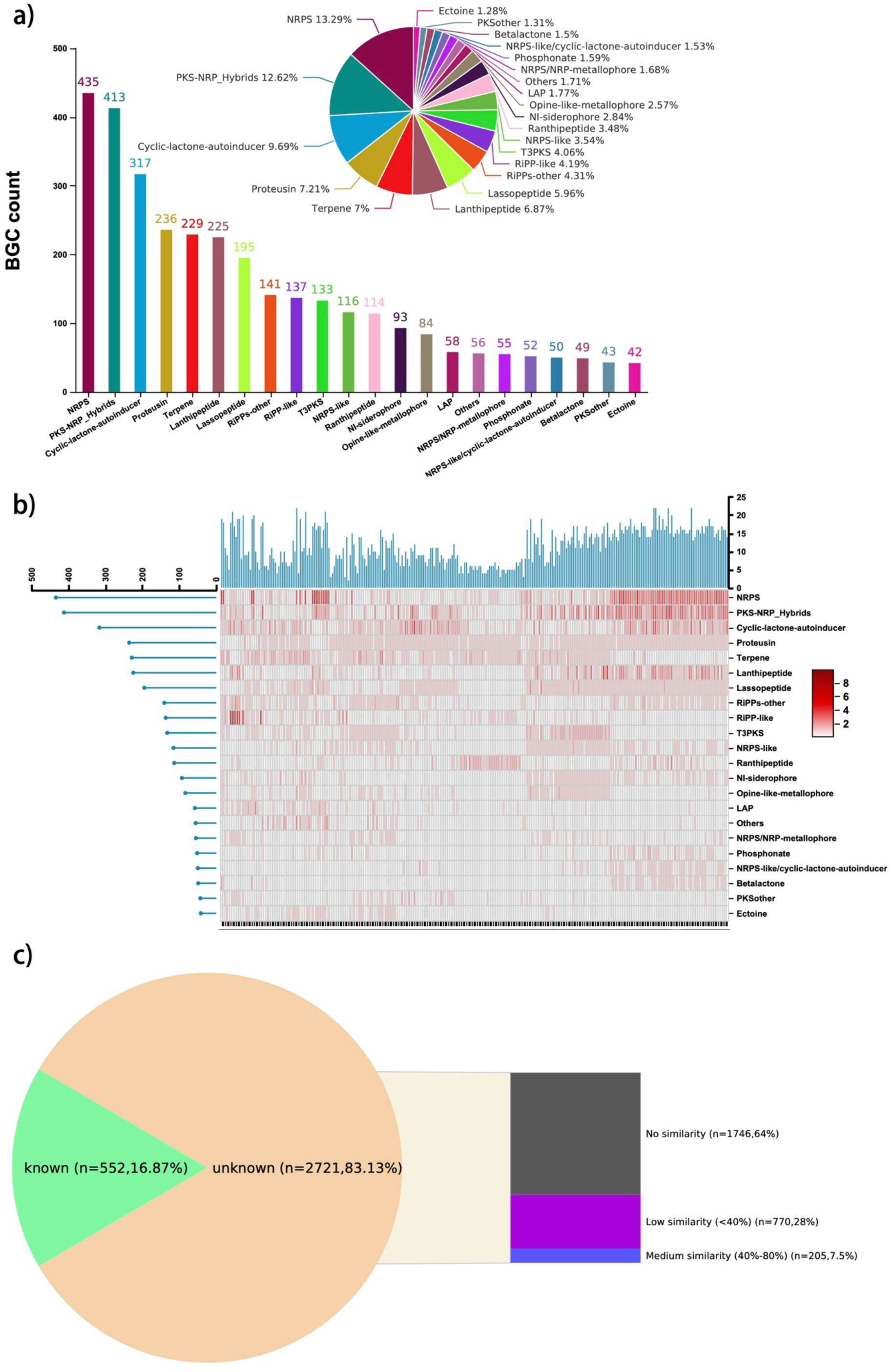
Biosynthetic gene clusters (BGCs) in the analyzed 284 genomes of the *Paenibacillus* genus. (a) Abundance of different BGC types. The x axis shows different BGC types, and the y axis shows the total number of each type of BGCs from the 284 genomes. The pie chart shows the proportion of different types of BGCs. (b) The distribution of each BGC type across all genomes was shown by the heatmap. Each row represents a BGC type, and each column represents a genome. The bar son the left represent the total numbers of each BGC type, and the bars on top represent the total numbers of BGCs identified in each genome. (c) The pie chart shows the number and percentage of known (similarity>80%) and unknown (similarity≤80%) BGCs according to antiSMASH KnownClusterBlast. The color-coded bar highlights the number and percentage of different degrees of similarity from unknown BGCs: gray indicates no similarity, purple indicates low similarity (less than 40%), and blue indicates medium similarity (between 40 and 80%).

To evaluate the potential of the detected BGCs to encode known secondary metabolites, we employed the “KnownClusterBlast” program within the antiSMASH analysis to estimate the similarity of the BGCs in comparison to the known BGCs of the MIBiG database version 3 (Minimum Information about a Biosynthetic Gene cluster) (36). From the 3,273 BGC, 552 (16.87%) gene clusters were classified as known with high similarity (>80%) to MIBIG entries, and 2,721 (83.13%) were unknown. This observation is consistent with what Kim *et al*. found in their study for 89 *Paenibacillus* genomes, in which 716 (84.4%) of the 848 BGCs they identified were classified as unknown, and the percentage is approximately 85-90% for the *Bacillus* genus (21, 37), whereas the percentage of unknown BGCs reported in the *Streptomyces* genus is 56.4% (38). These observations demonstrate the benefit of performing large genome-wide BGC studies in the *Paenibacillus* genus, where there is more as-yet untapped potential for encoding and synthesizing novel compounds. For the 2,721unknown BGCs, KnownClusterBlast hits 205 BGCs with medium similarity (40-80%), and 770 BGCs with low similarity (<40%) to MIBiG, and the other 1,746 BGCs do not have any hits to the MIBiG entries (Fig. 1c and Table S2).

In order to assess the diverse biosynthetic potential across the *Paenibacillus* genus, BGCs were clustered into gene cluster families (GCFs) by utilizing BiG-SCAPE (Biosynthetic Gene Similarity Clustering and Prospecting Engine) with 0.3 cutoff. BiG-SCAPE is designed to identify BGCs that are likely to produce similar or related secondary metabolites and to group BGCs into GCFs based on their sequence similarity and shared domain architecture (39). In this process, BiG-SCAPE also incorporates the MIBiG database, which provides information if a GCF is related to known BGCs. In total, 1,013 GCFs were clustered from the 3,273 BGCs predicted from 284 genomes. There are 697 GCFs consisting of only 1 BGC (singleton), this means that no other BGCs in the dataset or from the MIBiG database share a high enough level of similarity with this particular BGC to be grouped with it in the same family, suggesting they may produce distinct secondary metabolites or are from underexplored biosynthetic pathways. Additionally, the analysis revealed 266 GCFs with 2-10 BGCs, 29 GCFs with 11-30 BGCs, and a further 21 GCFs consisting of more than 30 BGCs (Fig. S3 and Table S3). With the aid of BIG-SCAPE, we could perform a similar comparison on GCFs with known BGCs in the MIBiG database and classify the GCFs as either known or unknown as well as return the hit BGC ID of MIBiG. In the 316 GCFs clustered from >1 BGC, only 10 were classified as known GCFs as they matched with at least one known BGC in the MIBiG database. The remaining 306 were classified as unknown, which are attributed to unknown or novel compounds. The BGC types of the 10 known GCFs are NRPS (3 GCFs), Lanthipeptide (3GCFs), PKS-NRPS hybrids (2 GCFs), Lassopeptide (1 GCF), and Opine-like-metallophore (1 GCF) (Fig. S3).

To inspect the distribution and enrichment of diverse GCFs across all genomes, we mapped GCFs that contain more than 30 BGCs to the phylogenetic tree of the 284 *Paenibacillus* spp (Fig. 2). The majority of these GCFs demonstrate clade-specific distribution patterns within the phylogenetic tree, except for GCF1, GCF4, and GCF7, which form separate clusters at disparate branches of the tree. Notably, no GCFs were widely distributed across all the genomes, which suggests the lack of uniformly conserved BGCs in the *Paenibacillus* genus. Unexpectedly, we also observed the absence of GCF in numerous clades across the phylogenetic tree, which may be attributed to the low degree of similarity among the BGCs within these clades. The clade-limited spread and absence of shared GCFs in certain clades reflect a broader diversity of BGCs across the whole genus and indicate the potential for the discovery of novel compounds, especially in the less characterized clades. Noteworthy, the frequently detected GCFs, which are only present in few genomes with very close phylogenetic branch distance, are most likely due to the high number of publicly available genomes of repetitively isolated species.

**Fig. 2.**
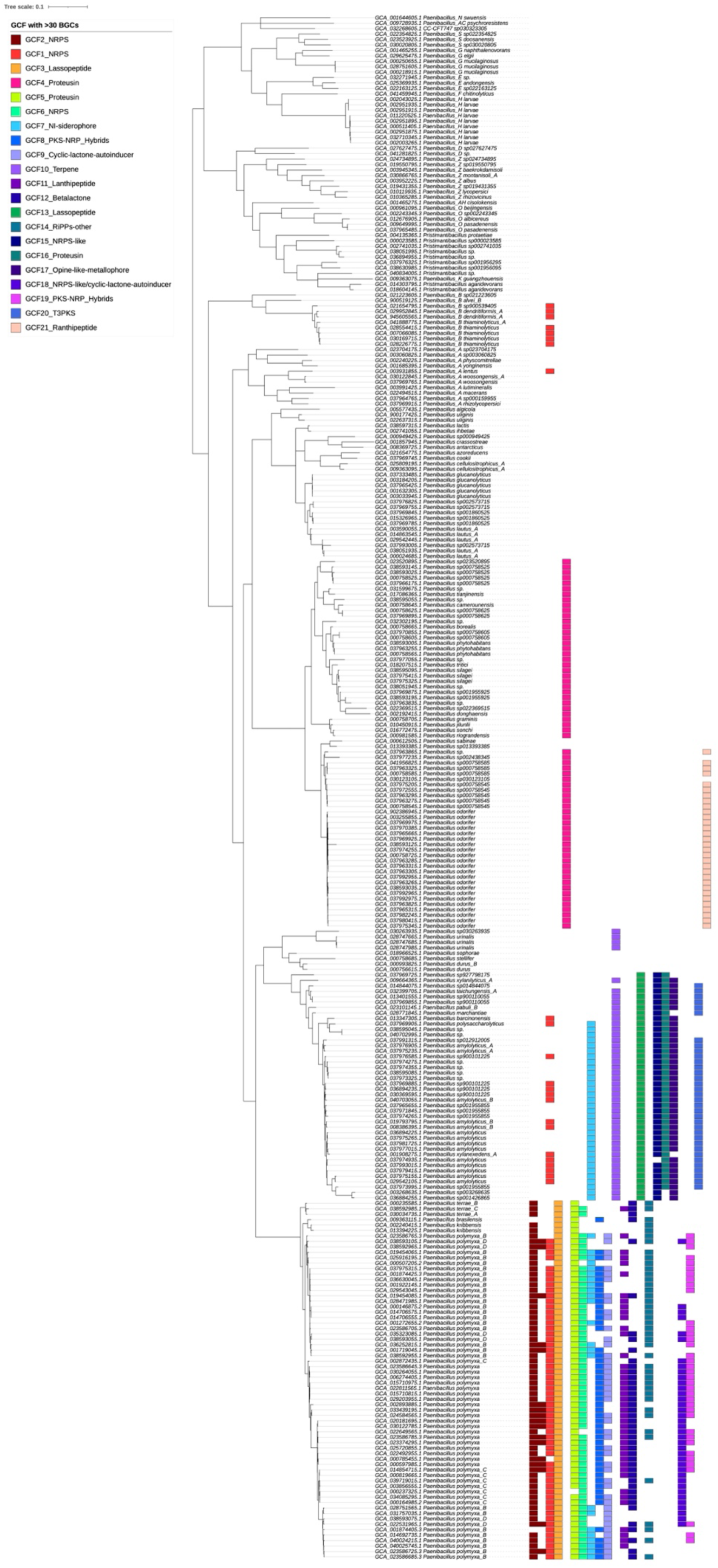
Phylogenetic tree and frequent GCF (with >30 BGCs detected) distribution of 284 *Paenibacillus* species.

### Genome dereplication reveals new patterns of abundance and distribution for BGC type within the *Paenibacillus* genus

To explore the true diversity and distribution of BGCs across different species within the *Paenibacillus* genus, we streamlined the dataset and minimized redundancy of the 284 genomes based on Mash distance, with a threshold of 0.05 (40). This strategy resulted in 126 representative genomes mostly representing one single species (Fig. S4 and Table S4). The detailed methodology is explained in the Methods section.

BGC mining, annotation, and statistical analysis were performed on the focused genome dataset (126 genomes) using the above-described methodologies. In total, 1,236 BGCs were identified in the non-redundant dataset, with the average of 9.81 BGCs per genome (see Table S5 and Fig. S5). This corresponds to previous studies on *Paenibacillus*, indicating that redundancy was adequately reduced in our focused dataset.

We compared the abundance and distribution of different types of BGCs in two datasets (the original 284 genomes and the 126 representative genomes after dereplication). In both datasets, the distribution and abundance patterns of major BGC types, such as NRPS, PKS-NRPS hybrids, cyclic-lactone autoinducer, proteusins, and the most abundant terpenes are all similar. This suggests that species of the *Paenibacillus* genus generally have the potential to synthesize these types of products. However, NRPS and PKS-NRPS hybrids were no longer the dominant two types of BGCs, but rather, cyclic-lactone-autoinducer, proteusins, and terpenes display comparable abundance (Fig. 3a). Matrix of BGC types and number against the phylogenetic tree of 126 genomes reveals that BGCs synthesizing terpenes and proteusins were the most widely distributed and were found in 93 genomes (73.81%). Other BGC types were identified in more than half of the genomes were cyclic-lactone-autoinducer (68.25%) and lassopeptide (59.25%) (Fig. 3b and Table 1). Although the total number of NRPS and PKS-NRPS hybrids types of BGCs was still high, they were identified in only 41.27% (52/126) and 46.83% (59/126) of the genomes, respectively, suggesting that they are enriched in some specific species. In the previous studies, NRPS was considered to represent the dominant BGC type (29, 30), but our extended study of all high-quality genomes within the *Paenibacillus* genus (after de-replication) reveals that this is not the case for many specific species or clades, such as species of *Paenibacillus_Z* clade (Fig. 3b).

**Fig. 3.**
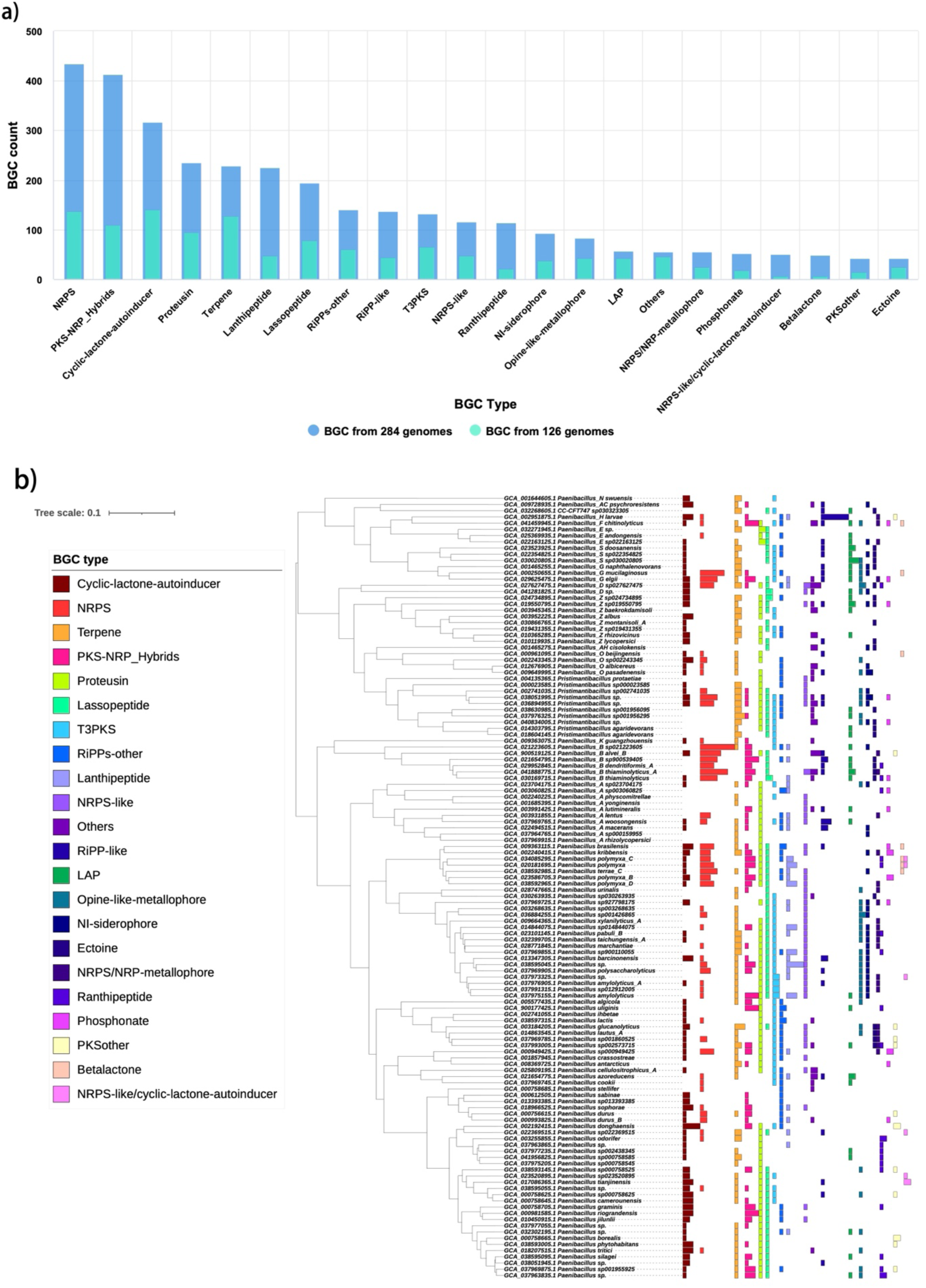
Abundances and distribution of BGCs in dataset of the 126 representative genomes. (a) Abundance of different BGC types in the two datasets. (b) Distribution of BGCs and BGC types across 126 representative isolates.

**Table 1.** The 1,236 BGCs from 126 representative genomes.

| **BGC type** | **BGC count** | **Genome count and coverage of 284 dataset** |  |
| --- | --- | --- | --- |
| Terpene | 127 | 93 | 73.81% |
| Proteusin | 95 | 93 | 73.81% |
| Cyclic-lactone-autoinducer | 141 | 86 | 68.25% |
| Lasso peptide | 78 | 75 | 59.52% |
| T3PKS | 65 | 61 | 48.41% |
| PKS-NRP_Hybrids | 110 | 59 | 46.83% |
| RiPPs-other | 60 | 56 | 44.44% |
| NRPS | 138 | 52 | 41.27% |
| NRPS-like | 48 | 46 | 36.51% |
| Opine-like-metallophore | 42 | 40 | 31.75% |
| LAP | 42 | 36 | 28.57% |
| NI-siderophore | 37 | 35 | 27.78% |
| Lanthipeptide | 48 | 33 | 26.19% |
| RiPP-like | 44 | 32 | 25.40% |
| Others | 45 | 31 | 24.60% |
| Ectoine | 24 | 24 | 19.05% |
| NRPS/NRP-metallophore | 24 | 24 | 19.05% |
| Ranthipeptide | 22 | 20 | 15.87% |
| Phosphonate | 18 | 15 | 11.90% |
| PKSother | 14 | 12 | 9.52% |
| Betalactone | 7 | 7 | 5.56% |
| NRPS-like/cyclic-lactone-autoinducer | 7 | 6 | 4.76% |

### The focused non-redundant genome dataset displays a greater diversity and novelty of secondary metabolite synthesis potential

The non-redundant representative genome dataset eliminates the interference caused by highly similar and redundant BGCs that are present in the same or very closely related species and therefore expected to provide a more unbiased information on BGC diversity. To better explore the synthetic capabilities of these BGCs, we use BIG-SCAPE to cluster BGCs into GCFs based on their similarity. From the 1,236 BGCs predicted from the 126 representative genomes, a total of 831 GCFs were identified, including 695 singletons, which accounted for 83.63% of all BGCs. Furthermore, 541 BGCs were clustered into 136 families, which included only 22 GCFs with more than 5 BGCs, highlighting that this dataset is non-redundant and the potential of BGCs is varied because similar BGCs will be clustered into GCF and are expected to synthesize the same or similar products. (Table S6 and Fig. 4).

**Fig. 4.**
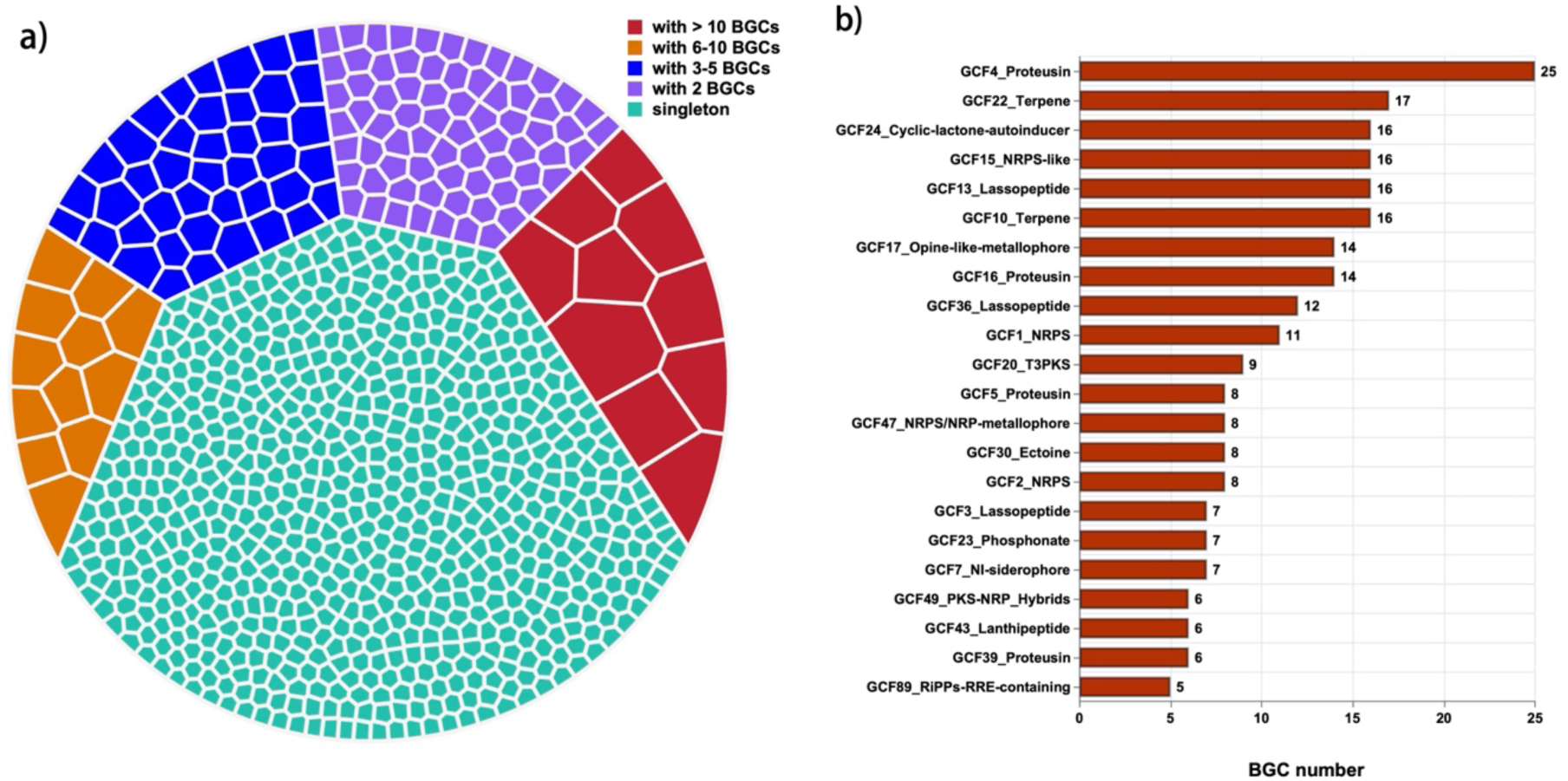
Depiction of the 831 GCFs clustered from the 1,236 BGCs predicted from 126 *Paenibacillus* genomes. (a) The color-coded pie chart illustrates the 831 GCFs with different size: green represents 695 singletons, purple represents 72 GCFs with 2 BGCs, blue represents 42 GCFs with 3-5 BGCs, orange represents 12 GCFs with 6-10 BGCs, and red represents 10 GCFs with more than 10 BGCs. Each small compartment represents a GCF, with the size representing the number of BGCs. (b) IDs and types of the 22 GCFs, which are each constituted by more than 5 BGCs.

Moreover, BIG-SCAPE incorporates known BGCs from the MIBiG database, thereby facilitating the identification of known and novel GCFs that encode previously uncharacterized products. Of the 831 GCFs, 11 (1.3%) were classified as known due to the presence of similar BGCs in the MIBiG database, including 5 singleton BGCs and 6 GCFs clustered from 4-11 BGCs (see Table 2).

**Table 2.**
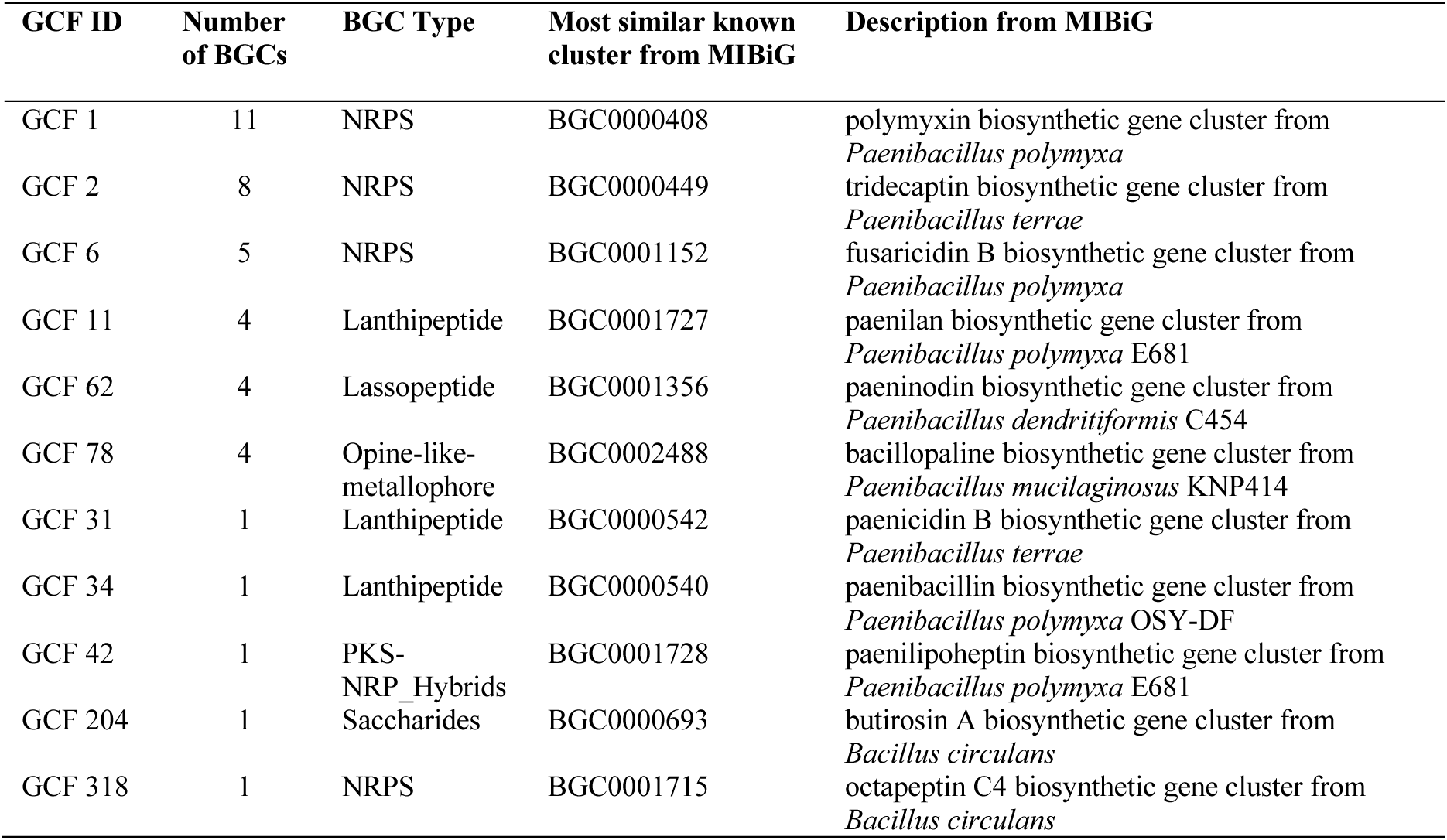
The 11 known GCFs of 126 representative genomes.

The top three “known” GCFs (CGF1, GCF2, and GCF 6) containing the highest number of BGCs are all of NRPS type, consisting of 11, 8, and 5 BGCs, respectively, and encode BGCs of the polymyxin, tridecaptin, and fusaricidin B-family, all of which are well-studied antibiotics produced by *Paenibacillus* spp (41–43). GCF 11, GCF 31, and GCF 34 all code for lanthipeptides, one of the most well-studied families of ribosomally synthesized and post-translationally modified peptides (RiPPs), and they are predicated to encode antimicrobial peptide produced by *Paenibacillus* spp, which are paenilan, paenicidin B, and paenibacillin (42, 44, 45). In summary, 9 of the 11 GCFs are related to synthesis of 9 known compounds in *Paenibacillus*, as recorded in the MIBiG database. Notably, although the known cluster most similar to GCF 204 was originally reported in *Bacillus circulans* SANK 72073, a recent study has suggested that this species should be reclassified to the *Paenibacillus* genus (46).

The remaining 820 unknown GCFs, which account for 98.7% of all GCFs, are of particular interest as they are likely to encode the biosynthesis apparatus of unknown secondary metabolites, and this promises exciting future discoveries. This includes 690 singletons, 20 relatively large families clustered by more than 5 BGCs, and other 110 GCFs.

### Phylogenetic distribution of GCFs across 126 representative genomes

To explore the biosynthetic potential of species across the *Paenibacillus* genus, we examined the phylogenetic distribution of large GCFs (comprising >5 BGCs) in 126 representative genomes (Fig. 5).

**Fig. 5.**
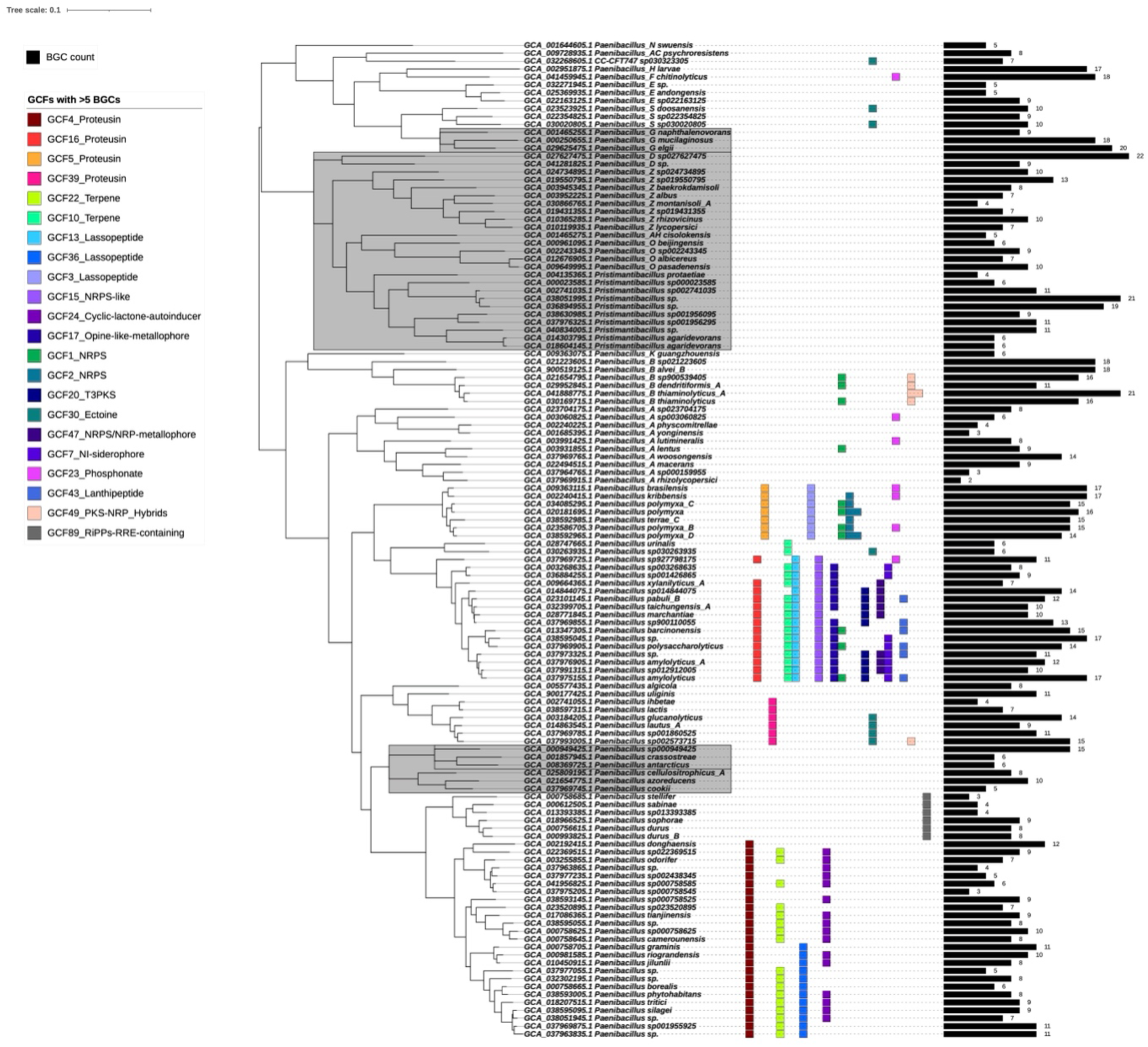
Phylogenetic distribution of large GCF (comprising >5 BGCs) in 126 representative genomes. The black bars represent the total number of BGCs in each genome. Each other small square represents a BGC, while squares of the same color represent a gene cluster family (GCFs).

Most GCFs demonstrated striking enrichment patterns that align with branching of the phylogenetic tree and correlated with the taxonomy depicted in the Genome Taxonomy Database (GTDB) release 09-RS220 (24th April 2024) (33), suggesting that these BGCs and their products are phylogenetically conserved and may reflect evolutionary conserved adaptations. The distinctively scattered GCFs and particular BGC products within phylogenetic clades and subclades indicates that these clades have developed unique biosynthetic capabilities during evolution to adapt to the ecological pressures and diversity of environmental ecological niches or these could be potentially horizontally transferred. GCF4 is a representative example, it clusters 25 BGCs of 25 genomes that is localized to a separate clade in the phylogenetic tree. GCF4 is found in all strains of this clade but absent in any other species. Despite the lack of comprehensive study and characterization of these proteusin-BGCs (47), the clade-specific distribution pattern provides a framework for further targeted exploration for its ecological role in these *Paenibacillus* species. Such clade specific GCF has been previously reported in the *Bacillus* (20) and *Streptomyces* (38) genera, and in our study in the *Paenibacillus* genus.

In addition, and more surprisingly, the complete absence of any conserved GCFs in some clades and subclades suggests that these species may have very distinct and diversified BGCs that have such low similarity to each other that they cannot be clustered into any GCFs. The lack of conserved GCFs again promises opportunities for discovering new or rare potential biosynthetic pathways in these taxa. Some of these clades that lack conserved GCFs are marked with a grey background in Figure 5, including species from *Paenibacillus_G*, *Paenibacillus_D*, *Paenibacillus_Z*, *Paenibacillus_O*, and some unclassified species. It is possible that they represent an important source of novel biologically active compounds with potential applications in biotechnology or pharmaceuticals, given the distinctive diversity and novelty of their biosynthetic gene clusters. However, it is also important to consider that the observed absence of conserved GCFs in these clades could be influenced by biases arising from the limited representation of genomes for certain species in the dataset. Inadequate sampling of these clades could lead to an underestimation of shared GCFs, as a larger genomic dataset might reveal conserved GCFs that are currently undetected.

## CONCLUSIONS

We performed a comprehensive gene mining for secondary metabolite biosynthetic gene clusters based on the full extent of available high-quality genomic data (as of October 2024) within the *Paenibacillus* genus. This has significantly expanded our understanding of the synthetic capacity of the species within this genus, and has led to the discovery of a highly diverse biosynthetic capacity for natural products. A large number of uncharacterized BGCs were identified and the majority of GCFs identified through BGC similarity networks could not be correlated with known compounds, which highlights the potential of *Paenibacillus* as a promising source for the discovery of novel compounds. We showed that most of identified GCFs are clade-specific and consistent with the phylogenetic tree after de-duplication, revealing that particular biosynthetic capabilities have evolved along specific branches of this genus over evolutionary time. Furthermore, species in other branches lacking GCFs are considered to have more diverse and novel BGCs, which makes them worthy of further attention and in-depth study.

## METHODS

### Dataset preparation

All available genomes of *Paenibacillus* genus were queried from NCBI in October 2024, and the genomes with ‘complete’ and ‘chromosome’ assembly levels designated at NCBI were selected. Subsequently, the quality of the genomes was evaluated using CheckM (version 1.2.2) (48), and high-quality genomes were chosen following a threshold of <5% for contamination and >90% for completeness and finally yielded the original dataset of 284 genomes. The analysis process is realized by BGCFlow, which integrates numerous software packages (7).

### Taxonomic classification and Tree building

To provide a more detailed description and exploration of the composition and distribution of secondary metabolite biosynthesis gene clusters across species in the *Paenibacillus* genus, we utilize GTDB-tk (version 2.4.0) and GTDB (release 09-RS220) (32, 33) for taxonomic classification and the definitions were used consistently throughout the data analysis and data visualization. The phylogenomic trees was constructed using the top 100 genes with the highest pre-calculated dN/dS values through a customized autoMLST wrapper which bypass additional organism selection (https://github.com/NBChub/automlst-simplified-wrapper) (20, 49). Tree visualizations were generated using Interactive Tree of Life (iTOL) (50).

### Genome mining analysis for BGCs

The process of genome mining for BGC is performed through the utilization of BGCFlow in the following steps (7). Annotated genomes were fetched from NCBI, after which the secondary metabolite BGCs were analyzed using the antiSMASH 7.1.0 (6). The antiSMASH KnownClusterBlast was employed to compare the similarity between the detected BGC regions and the characterized BGCs from the MIBiG 3.1 database (36) and to get a similarity level following this: similarity scores >80% is defined as high similarity, 40%-80% is defined as medium similarity, <40% is defined as low similarity, 0 score means no similarity.

### GCF clustering

We use BiG-SCAPE version 1.1.9 (39) with 0.3 cutoff for GCF clustering, which is based on the sequence similarity of all BGCs to identify which BGCs are likely to encode similar or the same products. These are then clustered into a family, while those BGCs that cannot be clustered with any other BGCs are defined as singletons. We further identified GCF as known if it has BGC with similarity above 80% against the MIBiG database calculated by *knownclusterblast*. Then the present and absent matrix was visualized in the phylogenetic tree using Interactive Tree of Life (iTOL).

### Genome de-replicating

To get a comprehensive and accurate representation of the diversity and distribution of BGCs across the species within the *Paenibacillus* genus, we obtained representative genomes in the initial dataset of 284 genomes by de-replication. The Mash distance between each pair of genomes was calculated. A threshold of Mash distance < 0.05 is used to identify which genomes are likely to be from the same species and cluster them into groups, then the best quality genome (higher completeness and lower contamination) is selected as the representative genome for each group, ensuring that the retained genomes were representative of their groups and of high quality, and others are removed as redundant.

## ACKNOWLEDGEMENTS

This project was supported by the Danish National Research Foundation (DNRF137) for the Center for Microbial Secondary Metabolites, and Novo Nordisk Foundation within the INTERACT project of the Collaborative Crop Resiliency Program (NNF19SA0059360). TW acknowledges funding from the Novo Nordisk Foundation Center for Biosustainability (NNF20CC0035580) and the Novo Nordisk Copenhagen Bioscience PhD program (NNF20SA0035588).

## Supplementary figures

Fig S1. Phylogenetic distribution and basic characters of 284 genomes from genus *Paenibacillus*.

Fig S2. BGCs identified from the 284 genomes.

Fig S3. The 1,013 GCFs clustered from the 3,273 BGCs predicted from 284 *Paenibacillus* genomes.

Fig S4. 126 representative genomes.

Fig S5. Biosynthetic gene clusters (BGC) in 126 representative genomes of genus *Paenibacillus*.

## Supplementary tables

Table S1. 284 genomes of *Paenibacillus*.

Table S2. The 3,273 BGCs identified from 284 genomes.

Table S3. The 1,013 GCFs clustered from the 3,273 BGCs predicted in 284 genomes.

Table S4. 126 representative genomes of genus *Paenibacillus*.

Table S5. The 1,236 BGCs identified from 126 representative genomes.

Table S6. The 831 GCFs clustered from the 1,236 BGCs predicted in 126 genomes.

