## Supplementary material for "Unlocking the biosynthetic potential of *Paenibacilli* through a genus-wide exploration of gene clusters for secondary metabolite production": Fig S1 to S5

### Supplementary Figures S1 to S5

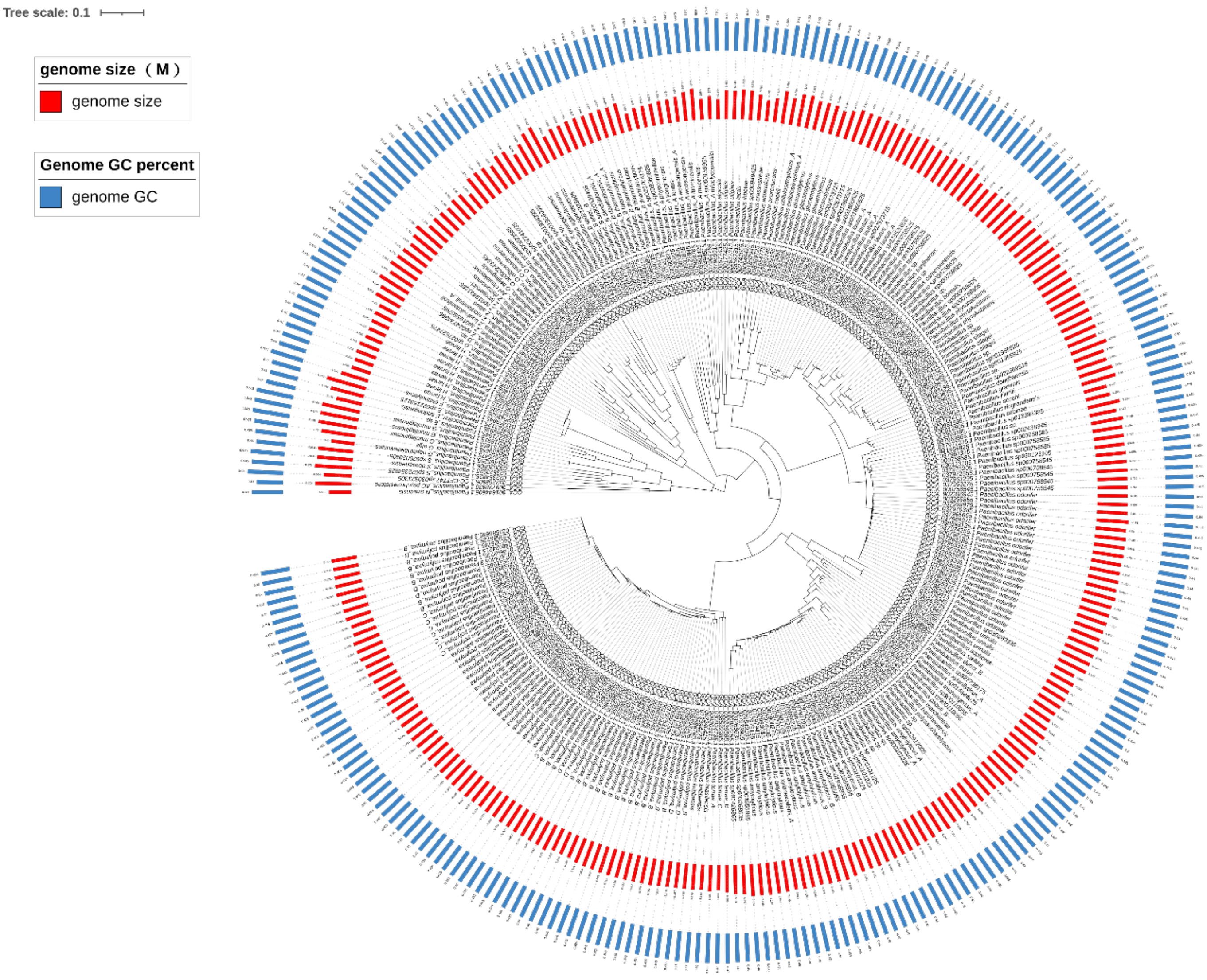


**Fig S1. Phylogenetic distribution and basic characters of 284 genomes from genus *Paenibacillus*.**
The phylogenetic tree was constructed based on multilocus sequence alignment (MLSA) of 30 genes and visualized in iTOL.

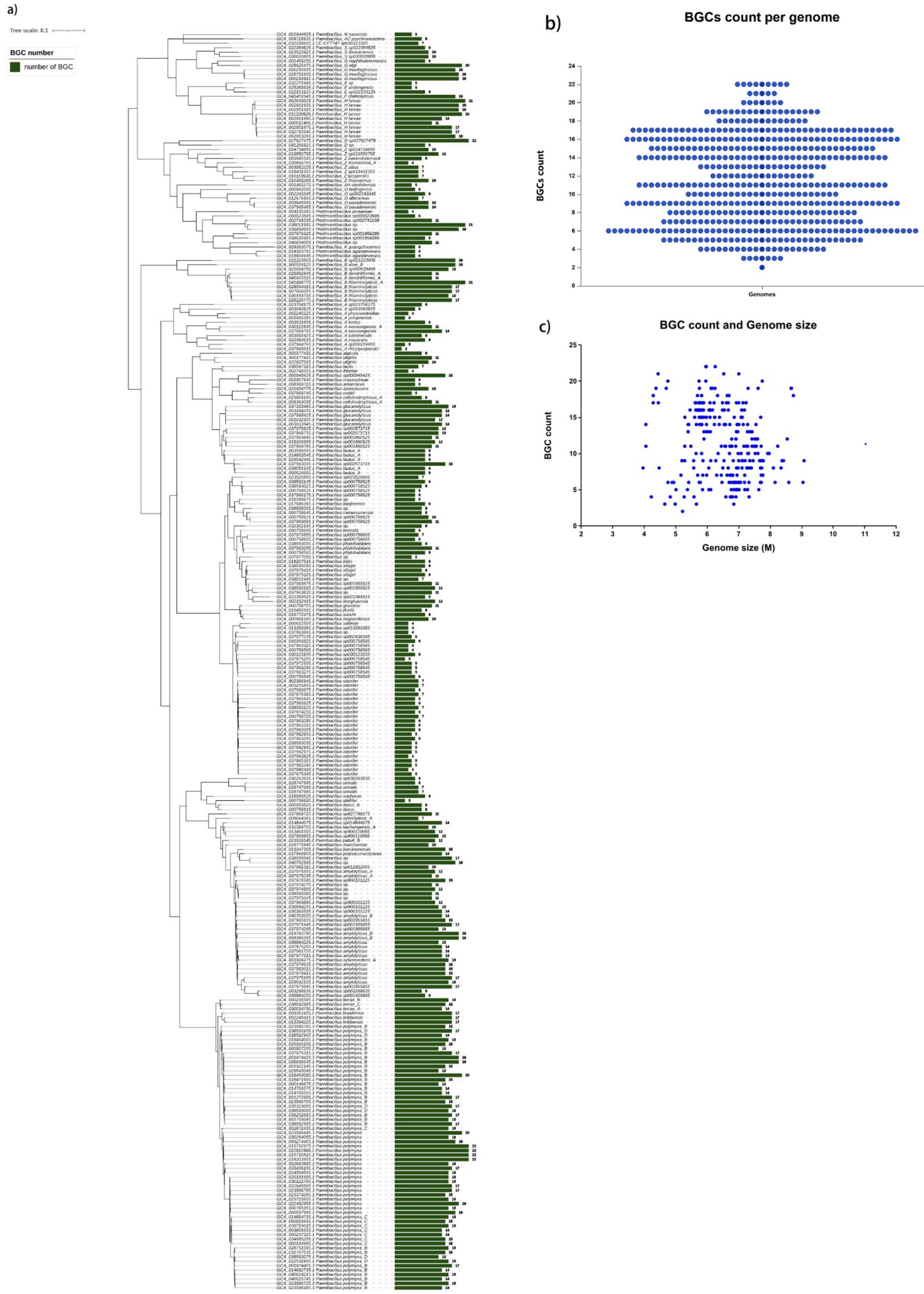


**Fig S2. BGCs identified from the 284 genomes.**

a) Phylogenetic tree was constructed based on MLSA. The red bar shows the BGCs number in each genome.

b) Overview of the abundance of BGCs across 284 genomes.

c) The relationship between genome size and the number of BGCs per genome. The x-axis is the genome size (Mbp), and the y-axis is the number of BGCs for each genome.

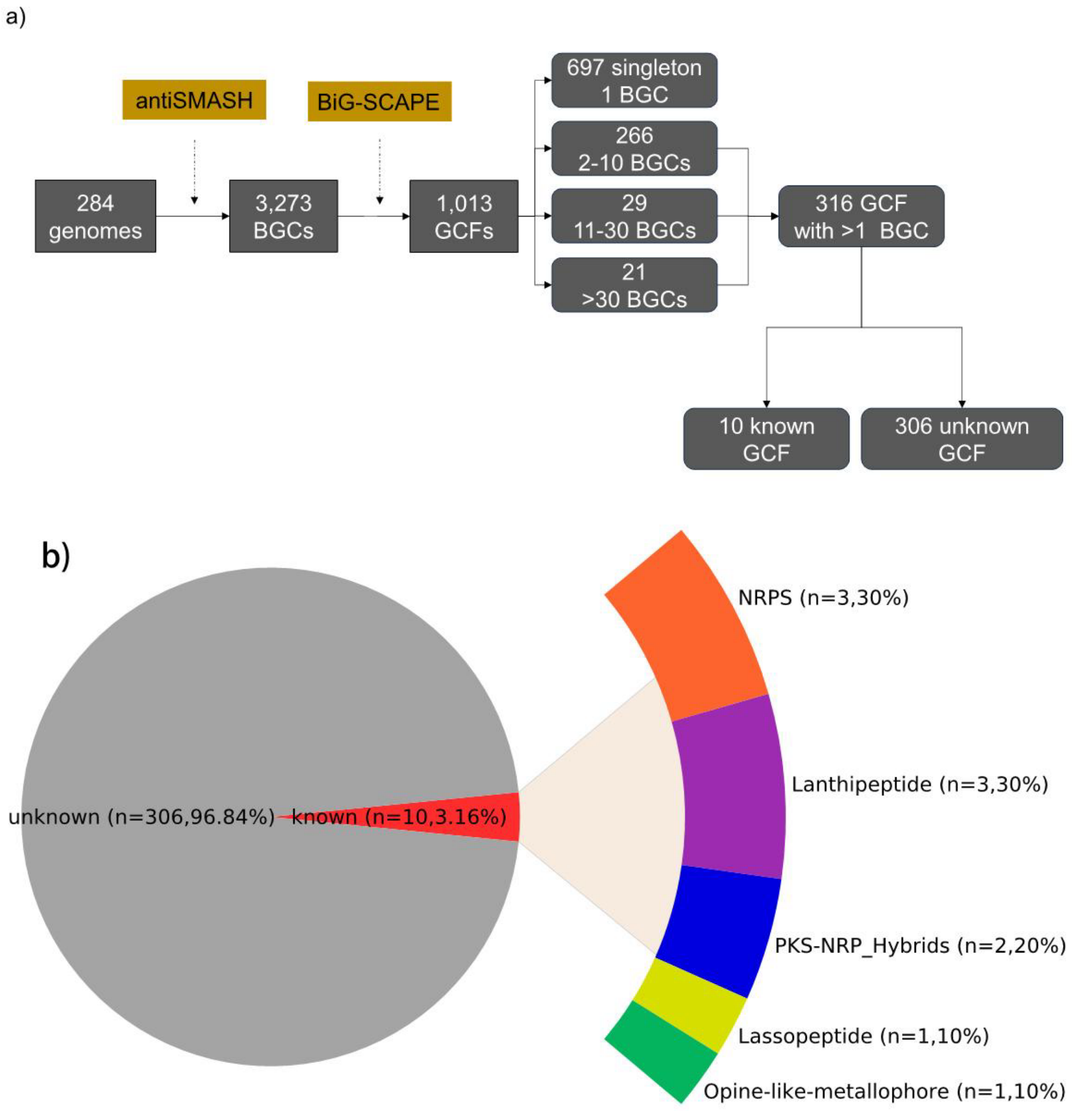


**Fig S3. The 1,013 GCFs clustered from the 3,273 BGCs predicted from 284 *Paenibacillus* genomes.**
a) Gene Cluster Families (GCFs) were clustered utilizing BiG-SCAPE, and known GCFs were identified by comparing with the MIBiG database.
b) The pie chart shows the number and percentage of known and unknown GCFs with more than 1 BGC. The color-coded bar indicates different types of the 10 known GCFs

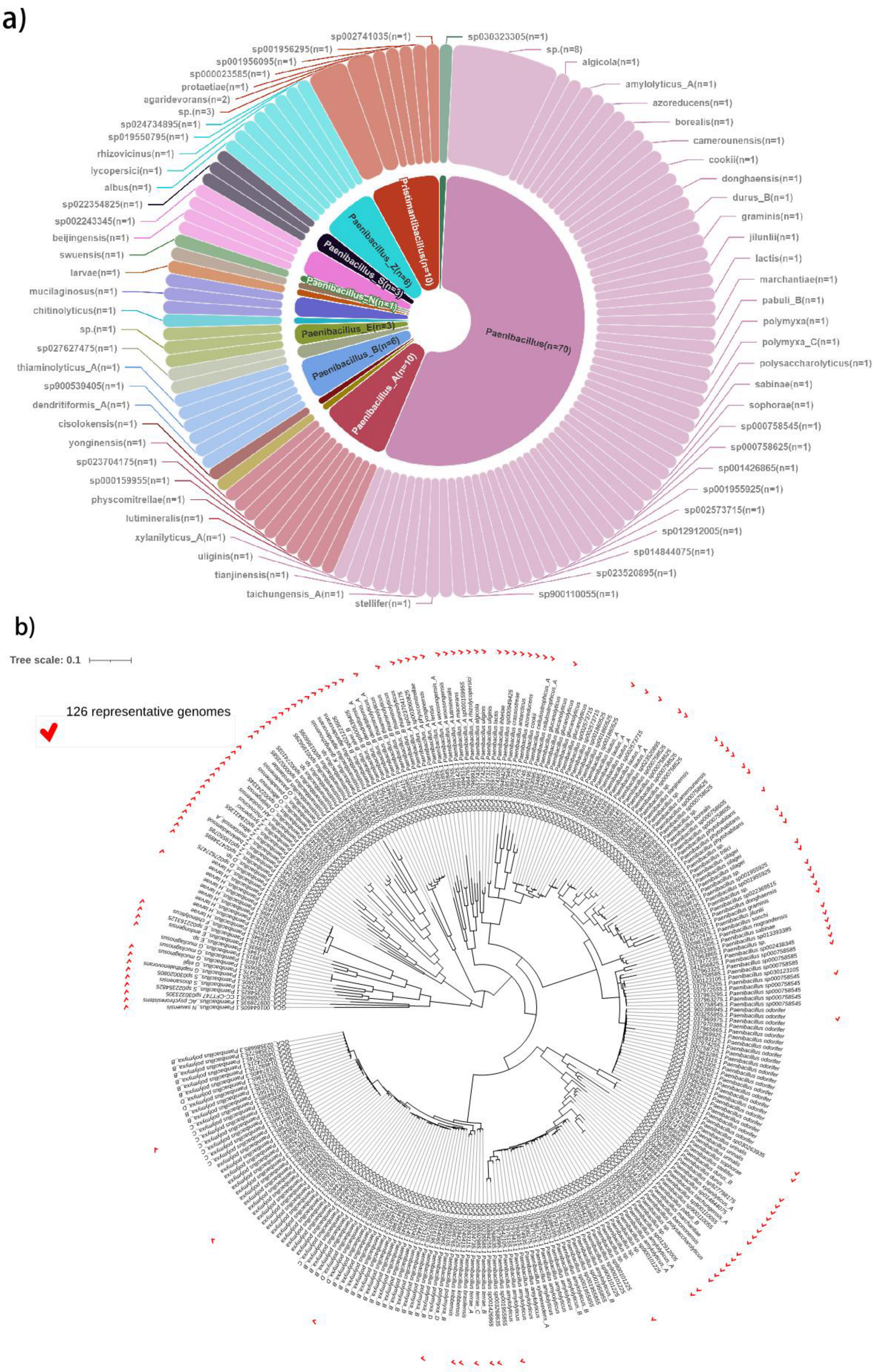


**Fig S4. 126 representative genomes.**

a) The circle diagram illustrates the species to which the 126 genomes belong, and most of the species were represented by a single genome.

b) 126 representative genomes were mapped to the phylogenetic tree of 284 genomes.

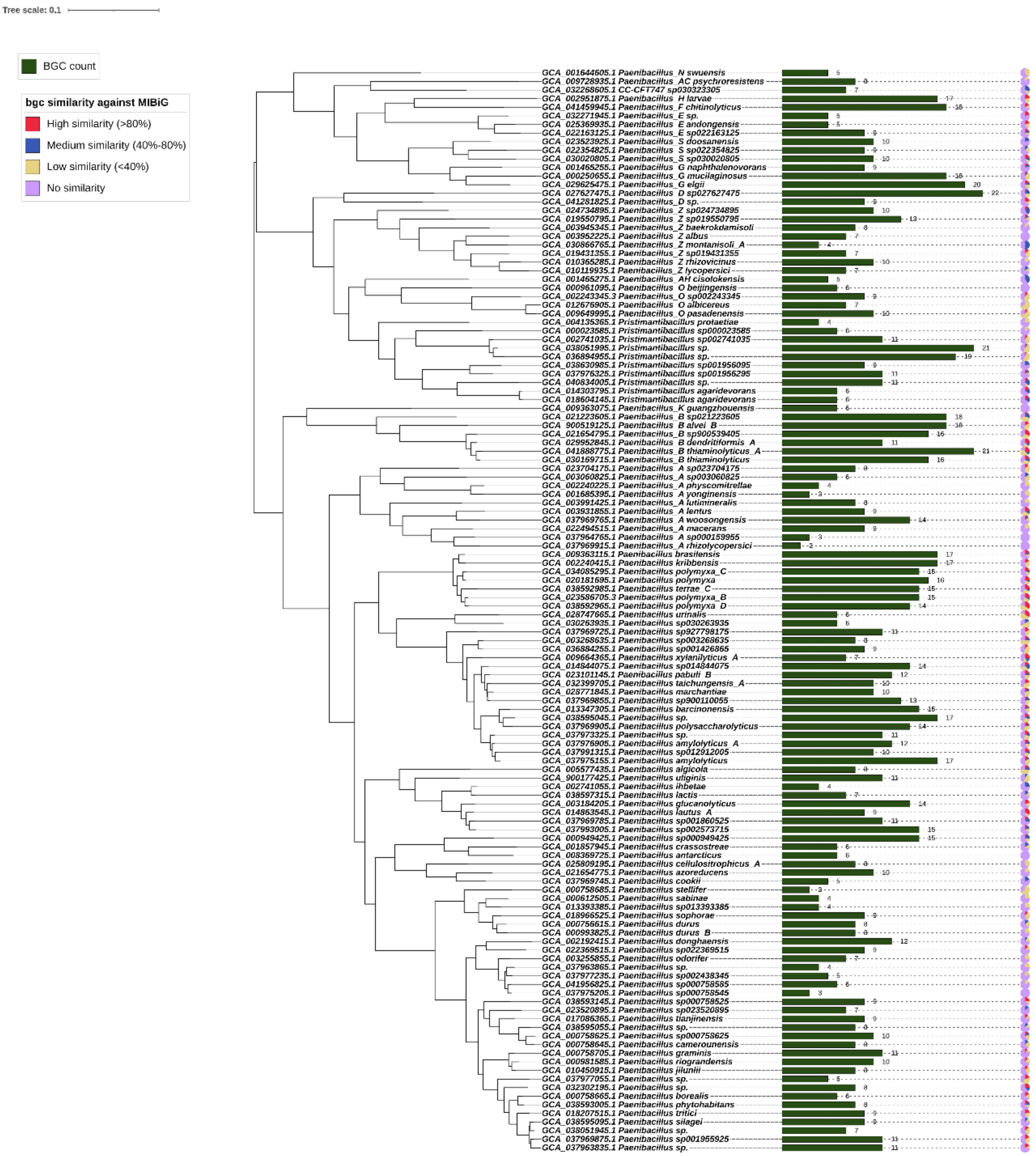


**Fig S5. Biosynthetic gene clusters (BGC) in 126 representative genomes of genus *Paenibacillus*.**

The green bar represents the number of BGCs in each genome, and the colored pie charts show the similarity of BGCs in each genome to known BGCs in the MIBiG database: purple indicates no similarity, yellow indicates low similarity (less than 40%), blue indicates medium similarity (between 40 and 80%), and red indicates high similarity (higher than 80%).
